# Modulating Bacterial Nanocellulose Crystallinity through Post-Transcriptional Repression in *Komagataeibacter xylinus*

**DOI:** 10.1101/2024.08.29.610269

**Authors:** Rahul Mangayil, Essi Sarlin, Tom Ellis, Ville Santala

## Abstract

Bacterial nanocellulose (BC), a versatile and biodegradable polymer, has been extensively studied as an alternative biomaterial for various applications. For biomedical and packaging uses, where precise control over nanocellulose structure is essential, existing literature describes BC structural modification processes that involve additives or additional steps. With the aim to develop a programmable method to control the bulk microstructure, we developed sRNA-based post-transcriptional repression cassettes that allows precise regulation of the crystalline phase of BC. Before investigating the effects of post-transcriptional repression of *bcsD, bcsZ*, and *ccpA* genes, known to influence BC crystallinity, we validated the specificity of the sRNA repression cassette by targeting a genome-integrated red fluorescent protein, mRFP1. The observed growth inhibition (>80%) caused by overexpressed Hfq RNA chaperone in *Komagataeibacter xylinus* was alleviated (≤ 23%) by its removal, resulting in a 43% reduction in relative mRFP1 expression. By varying the design of the repression cassette and the inducer concentrations, we successfully modulated the repression of the target genes [with relative expression reductions of 6%-34% for bcsD, 8%-24% for bcsZ, and 2%-20% for ccpA, as confirmed by qRT-PCR]. These gene repression levels led to statistically significant changes in the amorphous content of the BC microstructure, as demonstrated by X-ray diffraction and wide-angle X-ray scattering analysis.

## Introduction

Bacterial nanocellulose (BC) is as a versatile biopolymer with remarkable qualities, such as purity (devoid of hemicellulose and lignin), exceptional structural integrity, biodegradability, and water retention capabilities. Extensive research has delved into its applications, particularly in the biomedical and packaging sectors, where precise control over nanocellulose structure is paramount. Modifications to BC structures have predominantly relied on two approaches: in situ, by supplementing growth medium with additives (such as carboxymethyl cellulose and polyethylene glycol) (de Lima Fontes et al., 2018; Heßler & Klemm, 2009), and ex situ methods involving acetylation (Ávila Ramírez et al., 2016), phosphorylation (Busuioc et al., 2022), and physical processing (Aditya et al., 2022). However, such approaches often lead to bioprocess restrictions, additional downstream steps, and limits in situ control owing to the high hygroscopic nature of BC and the randomly orientated nanofibrils (Cheung et al., 2023; Murugarren et al., 2022)

To overcome these challenges, a comprehensive understanding of the genetics underpinning nanocellulose production in *Komagataeibacter* spp. is imperative. It is well known that bcs (acs, another acronym in literature) operon is the machinery involved in cellulose biogenesis (Romling & Galperin, 2015). Though a rational engineering target, lack of specific tools has limited genetic exploitations of *Komagataeibacter* bcs machinery, directing scientists to conduct heterologous expression of *Komagataeibacter* bcs operon in model bacterium such as *Escherichia coli* (Buldum et al., 2018; Buldum et al., 2020). However, the publications by Florea et al. (2016) and Teh et al. (2018) have opened genetic manipulation prospects in *Komagataeibacter* spp. (Florea et al., 2016; Mangayil et al., 2017; Teh et al., 2019)

BC synthesis is a biochemically complex process (Abidi et al., 2022). In *Komagataeibacter* spp., bcs operon comprising bcsAB (catalytic and regulatory genes for the polymerization of linear glucan chains), bcsC (involved in glucan chain secretion), and bcsD (responsible for fibril packing and conferring crystallinity), serves as the biogenetic machinery for cellulose synthesis (Romling & Galperin, 2015; Sana et al., 2024). In a study by Saxena et al. (1994), the functional roles of the bcs operon genes were individually assessed using a gene knockout (KO) approach. Their study found that mutants lacking the *bcsABC* genes failed to produce the biopolymer, underscoring the critical role of these genes in BC synthesis. In contrast, *bcsD* mutants deficient produced cellulose in lower yields with loosely packed structures, highlighting the importance of bcsD in nanocellulose assembly. This finding was corroborated in Hu et al. (2010) study, wherein the authors observed an association of bcsD with bcsC in BC secretion (Hu et al., 2010). Morphological, and structural changes through partial and complete overexpression of *K. xylinus* bcs operon was observed in our previous work (Mangayil et al., 2017). Compared to the BC produced by wildtype (WT) *Komagataeibacter*, overexpressed bcs operon (*bcsABCD*) led to improved production metrics, with the resulting biomaterial exhibiting densely packed cellulose nanofibrils. In contrast, partial overexpression of the bcs operon (*bcsA* and *bcsAB*) resulted in less intact BC.

In *Komagataeibacter* spp., the role of bcsD to partake alone in cellulose crystallization is argumentative. *Komagataeibacter* genome contains accessory genes, *bcsZ* (encoding for beta-1,4-endoglucanase, also reported as *CMCax* in literature), *ccpA* (cellulose complementing factor, also reported as *ccpAx* in literature), and *bglX* (encoding for glucosidase), upstream and downstream of bcs operon, respectively (Cannazza et al., 2021; Cannazza et al., 2022; Mangayil et al., 2020). Nakai et al. (2002) highlighted the positive impact of *ccpA* gene on BC crystallinity (Nakai et al., 2002; Sunagawa et al., 2013). Furthermore, ccpA has been speculated to closely associate with bcsD for effective localization of the bcs terminal complex (Kondo et al., 2022; Sana et al., 2024). Further investigations by Nakai et al. (2013) into the role of bcs operon accessory genes in BC biogenesis revealed that *K. xylinus* strains with a disrupted *bcsZ* gene to produce particulate nanocellulose with significant structural differences (Nakai et al., 2013). Although not directly related to *K. xylinus*, a study by Sajadi et al. (2019) on a native cellulose-producing *E. coli* strain provides some insights into the association between bcsD and BC regulatory proteins (Sajadi et al., 2019). The cellulose synthase machinery (*bcsABZC*) in *E. coli* Nissile 1917 lacks *bcsD*. However, upon *bcsD* overexpression, the authors observed decreased amorphous regions and improved crystallinity, suggesting complementary interactions between the native bcsZ and the heterologously expressed bcsD.

Diverging from the KO and overexpression methods employed in aforementioned studies, here we introduce a post-transcriptional repression system based on synthetic regulatory small RNAs (sRNA) to explore its suitability in *K. xylinus* and study its impact of post-transcriptional modulation of bcsD, bcsZ, and ccpA on BC crystallinity.

## Materials and Methods

### Bacterial strain and cultivation conditions

The cloning host, *Escherichia coli* XL1 (Stratagene, USA) was cultivated in low salt Lysogeny Broth medium (LB medium, pH 7.0) [g L^−1^: tryptone, 10; yeast extract, 5; sodium chloride, 5] supplemented with glucose (4 g L^−1^) at 37°C and 300 rpm. For recombinant strains, chloramphenicol (Cm; 34 μg ml^−1^) was supplemented to maintain the plasmid.

*K. xylinus* DSM 2325 (DSMZ, Germany) was the BC producing strain used in this study. For BC production, *K. xylinus* was grown statically in 6-well plates (Argos Technologies, Cole-Parmer, US) containing Hestrin–Schramm medium (HS medium, pH 6.0) [g L^−1^: peptone, 5; yeast extract, 5; disodium hydrogen phosphate, 2.7 and citric acid, 1.15) supplemented with glucose (20 g L^−1^). Chloramphenicol (340 μg ml^−1^) was supplemented to recombinant *K. xylinus* cultures to maintain the plasmid. Electrocompetent cells were prepared and transformation was conducted as described in our previous work (Mangayil et al., 2017). Precultivation of *K. xylinus* strains were conducted statically in HS medium supplemented with 2% glucose and 340 μg ml−1 ml^−1^ Cm (for recombinant strains). After 5 days of static cultivation, the BC pellicles were lysed overnight with 2% cellulase to obtain the entrapped cells, which were washed in sterile 1XPBS and resuspended in the same.

### Plasmid construction

The primer sequences used in this study are presented in Table 1.

**Table 1.**
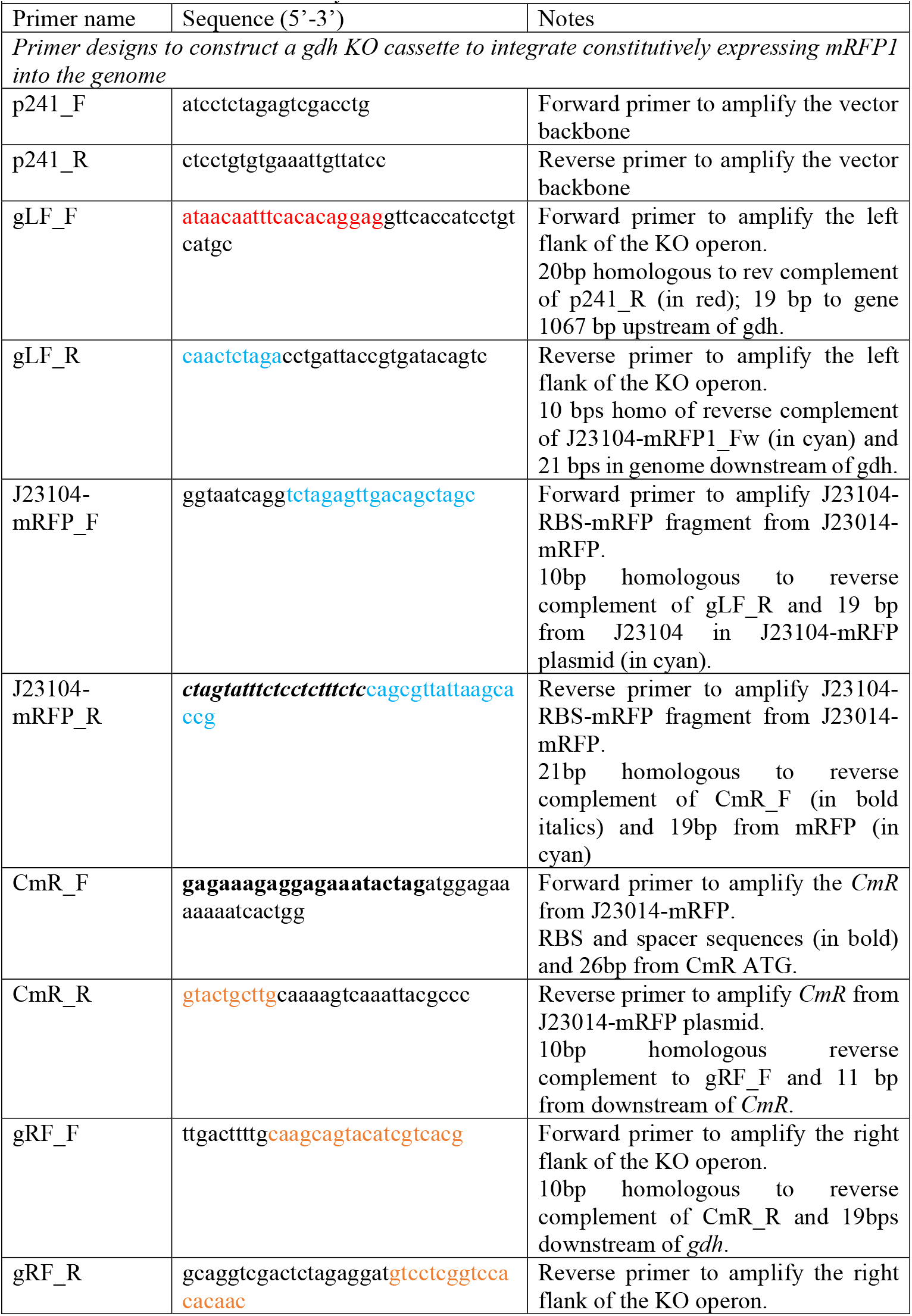
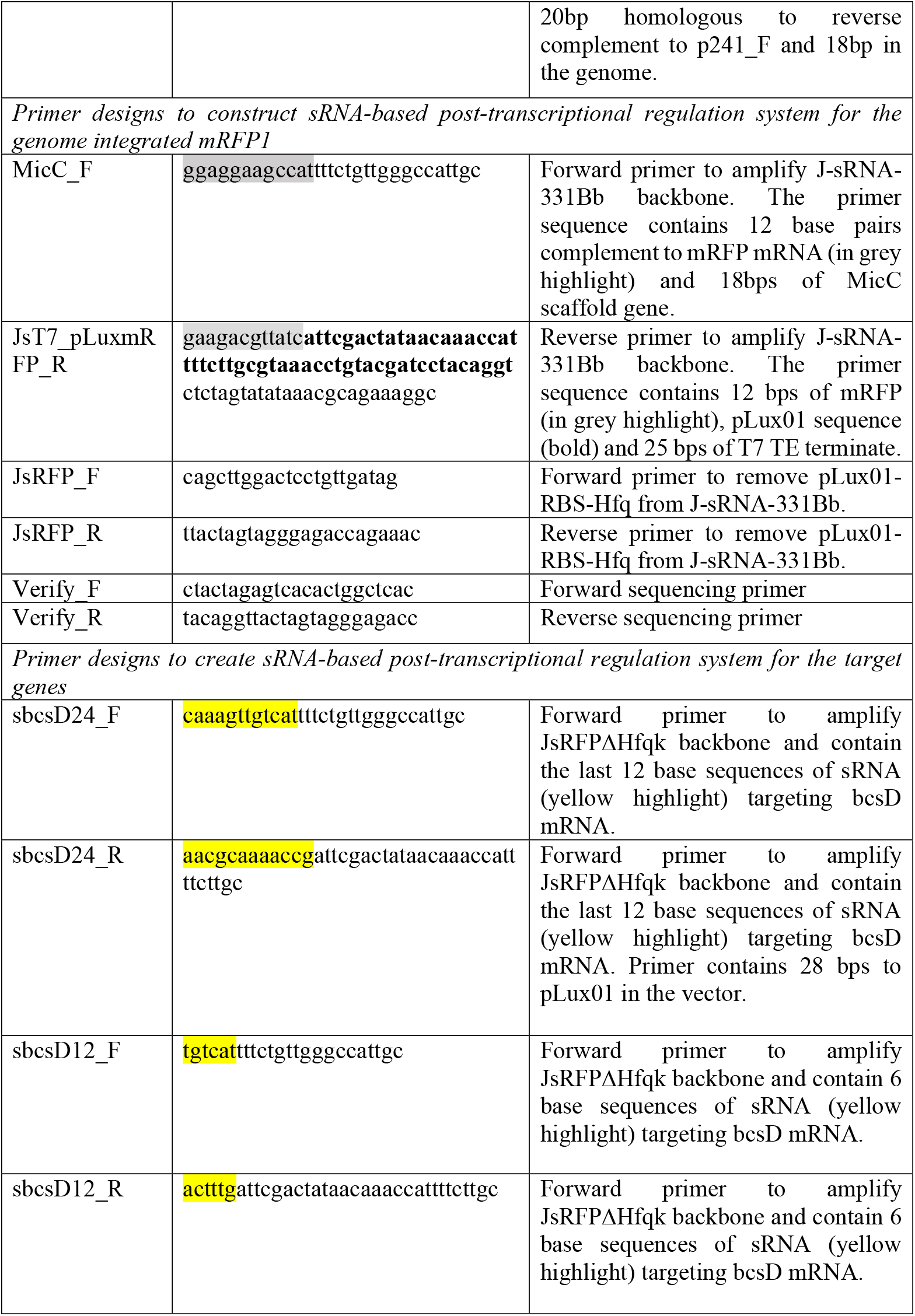
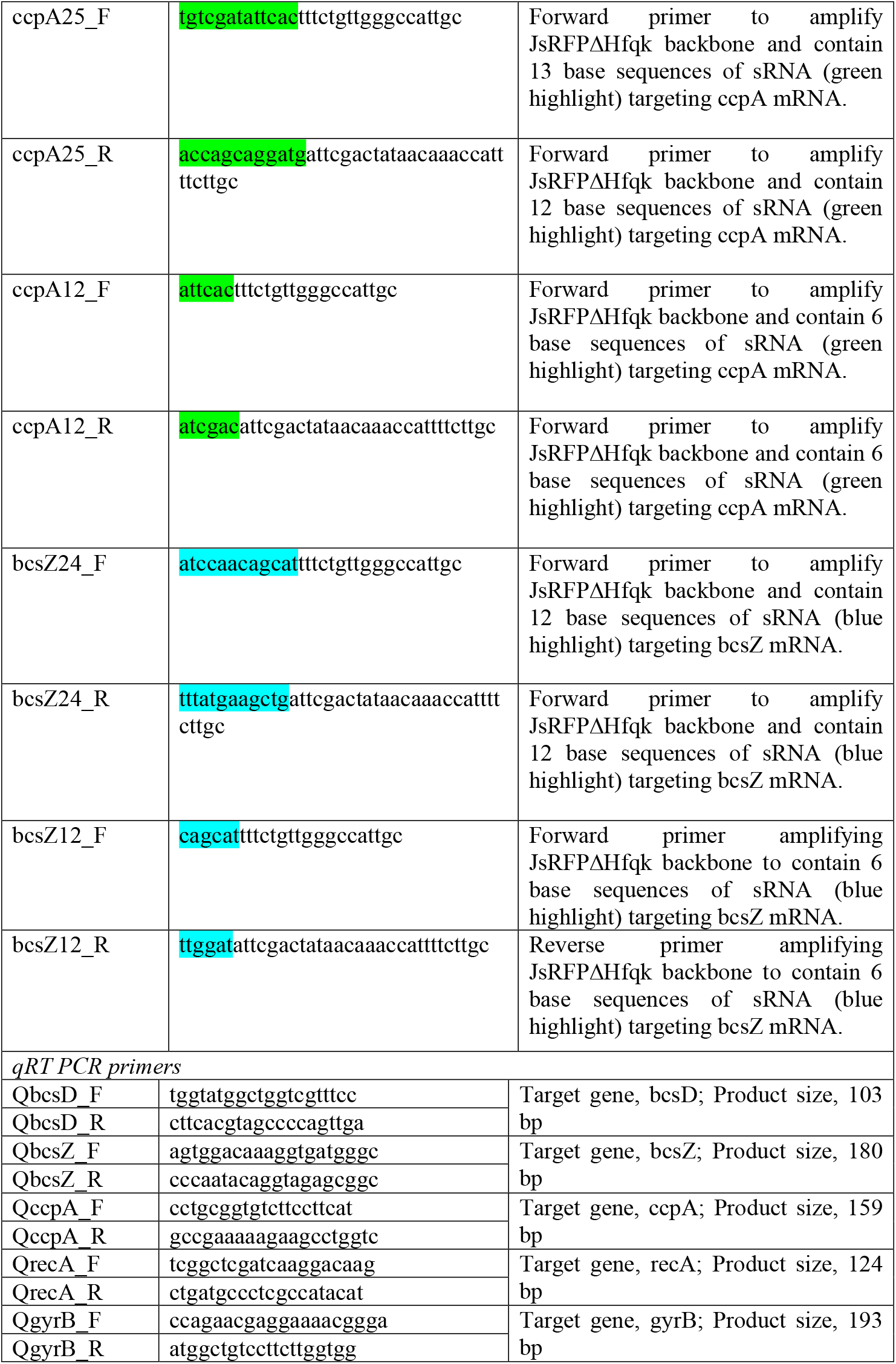
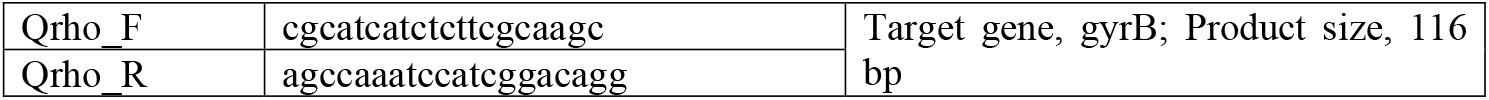
Primers used in this study.

The J23104-mRFP1 plasmid (Florea et al., 2016), which drives mRFP1 expression under a strong constitutive promoter, along with pSEVA241, kindly provided by the SEVA database curators upon request (https://seva-plasmids.com/), were used to design the genome-integrated, constitutively expressed mRFP1. The J-sRNA-331Bb plasmid [containing *E. coli* Hfq, sRNA targeting UDP-glucose pyrophosphorylase mRNA, and *E. coli* MicC scaffold under an acyl homoserine lactone (AHL)-inducible promoter] was used to design the post-transcriptional repression cassettes in this study (Florea et al., 2016).

The preliminary validation of post-transcriptional regulation construct was conducted using constitutively expressed *mRFP1* gene. Since the study targets *K. xylinus* genes, we decided to integrate the mRFP1 expression operon into the genome via homologous recombination. The glucose dehydrogenase gene (gdh, located at genome position 814017-816407 bp) was chosen as the recombination site (C. H. Kuo et al., 2015). For the same, ∼1kb fragments from sequences 1067 bp upstream (gLF) and 7 bp downstream of *gdh* (gRF) were amplified from the genome using primers gLF_F/gLF_R and gRF_F/gRF_R, respectively. The J23104-RBS-mRFP1 (JmRFP1) and RBS-chloramphenicol resistance (CmR) fragments from J23104-mRFP1 plasmid were amplified using primers J23104-mRFP_F/J23104-mRFP_R and CmR_Xyl_F/CmR_R, respectively. The *gdh* knock-out (KO) cassette, of design gLF-JmRFP1-CmR-gRF, was assembled into non-replicative pSEVA241 plasmid using Gibson assembly in *E. coli*. Following the validations, the construct was transformed into *K. xylinus* and the transformants (*K. xylinus* Δ*gdh*) were selected in HS agar containing 20 g L^-1^ glucose, 340 μg ml^-1^ Cm and 2% cellulase. Subsequently, fluorescent colonies were picked and confirmed by colony PCR.

### sRNA design

The sRNA-based post-transcriptional regulation system for the genome integrated *mRFP1* in *K. xylinus* Δ*gdh* was designed in J-sRNA-331Bb (using primers MicC_F and JsT7_pLuxmRFP_R). This system was intended to replace the existing sRNA, which targets the UDP-glucose pyrophosphorylase gene. The sRNAs were designed based on the protocol published by Yoo et al. (2013) (Yoo et al., 2013). sRNA sequences with binding efficiencies greater than −30 kcal mol^−1^ were selected to construct the repression plasmids and the mRNA binding efficiencies of the sRNA sequences were analyzed in silico using IntaRNA(Mann et al., 2017). Primers were designed to contain the sRNA sequences targeting mRFP1 mRNA are presented in Table 1. The linear PCR product containing the vector backbone, genetic elements and the sRNA sequences were recircularized using T4 PNK (Thermo Fischer Scientific, USA) and T4 ligase (Thermo Fischer Scientific, USA), resulting in JsRFP plasmid. As both JsRFP plasmid and the mRFP1 integration cassette contains *CmR*, the antibiotic resistance gene in JsRFP was replaced with kanamycin resistance gene (*kanR*) using SwaI and PshA restriction sites, resulting in JsRFPk (Silva-Rocha et al., 2013). To construct a version without the pLux01-RBS-Hfq operon, the JsRFPk vector was amplified using primers JsRFP_F and JsRFP_R and recircularized as mentioned previously, resulting in JsRFPΔHfqk plasmid.

Similar considerations were employed to design sRNAs targeting bcsD, ccpA, and bcsZ mRNAs (Table 1). To explore the potential for tunable post-transcriptional repression, shorter sRNAs consisting of 12 base pairs were designed. The primer pairs incorporating these sRNA sequences for each target gene (designated with ‘*gene name*’ and ‘number of sRNA base sequences’), along with their predicted binding energies, are detailed in Table 2. To achieve controlled repression of the target bcs genes, the sRNA sequence in JsRFPΔHfqk was replaced with sRNA sequences for the target genes (as detailed in Table 1 and Table 2), and the PCR product was recircularized. Upon verifying the constructs by sequencing (Macrogen, Netherlands) using the verify_F and verify_R primers, the plasmids were transformed into *K. xylinus*. The *K. xylinus* constructs harboring the repression plasmids with sRNA sequences targeting bcsD, ccpA, and bcsZ mRNAs are designated with ‘*gene name*’ and ‘number of sRNA base sequences’.

**Table 2.**
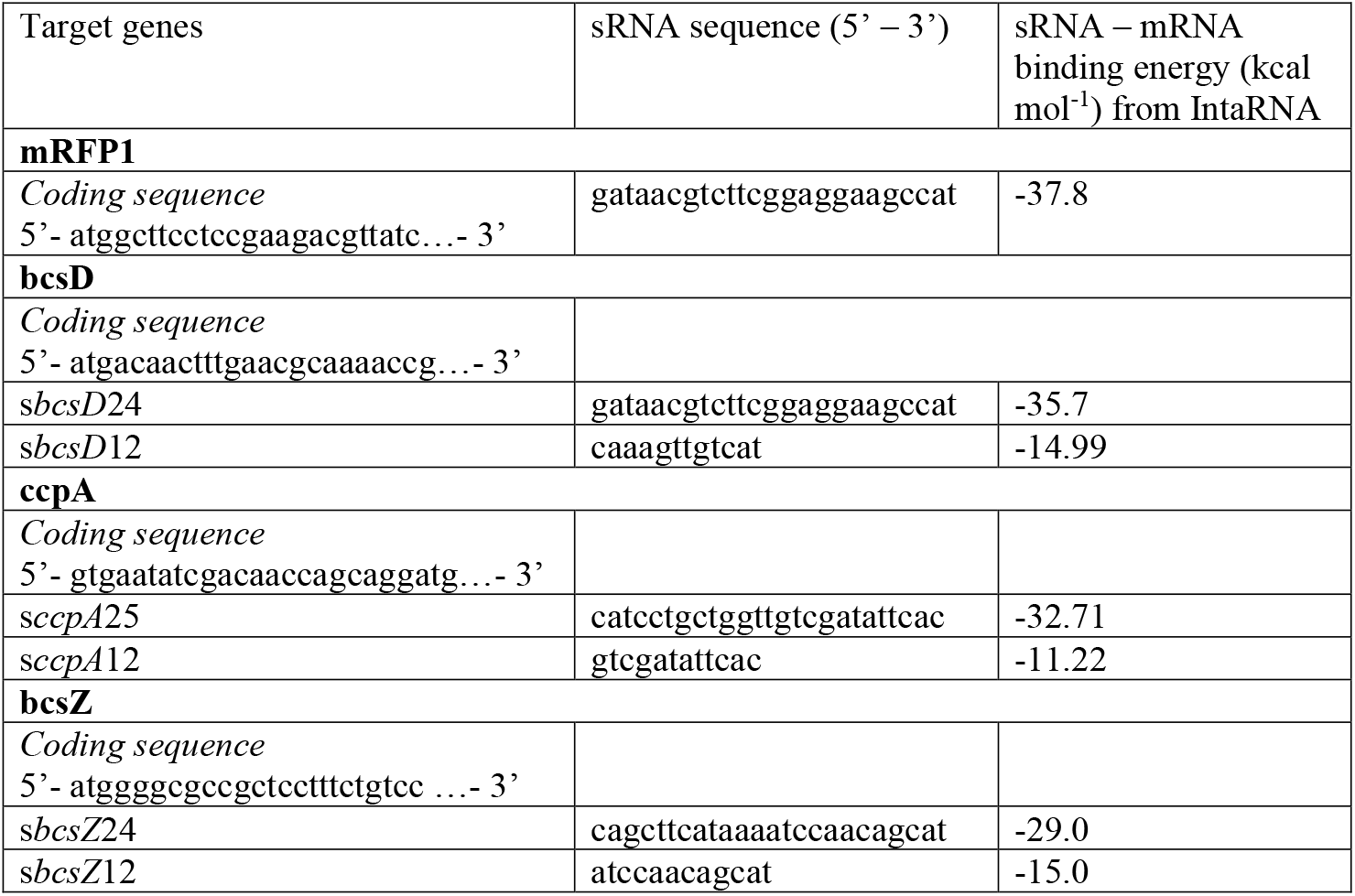
Coding sequence, designed sRNA sequence and the sRNA – mRNA binding energy predicted by IntaRNA.

### Fluorescence measurements

The WT and recombinant *K. xylinus* cells (*Kx*Δ*gdh*-JsRFPk and *K. xylinus*-J23104-mRFP1) harboring constitutively expressed mRFP1 in the genome and in plasmid, respectively, were inoculated with an initial cell density of 0.04 OD600nm in HS medium containing 2% glucose, 300 μg/ml Cm, 2% cellulase, was grown at 30°C and 180 rpm. Two hundred microliter of washed cells were pipetted to micropipette plate wells in triplicates and the fluorescence measurements were conducted at excitation and emission wavelengths of 584nm and 612nm, respectively, in Spark microplate reader (Tecan, Switzerland). To study the effect of Hfq overexpression in *Kx*Δ*gdh* cells on cell growth (OD_600nm_) and RFP expression, AHL induced and noninduced culture samples were collected every 24 hours.

### Quantitative real-time PCR

The qPCR primers (Table 1) were designed in Primer-Blast using respective NCBI GenBank accession numbers of the target genes (*bcsD*, WP_061271618.1; *bcsZ*, WP_007399068.1 and *ccpA*, WP_010513891.1). The primers were designed with a melting temperature of 60°C, a GC content of 55%, and no more than three consecutive G or C bases, aiming to generate amplicons of 100–200 bp. The primer sequences were analyzed using IDT DNA Oligo Analyzer tool (https://www.idtdna.com/pages/tools/oligoanalyzer) to predict secondary structures and primer-dimers. Additionally, the specificity of the primers for their target sequences was manually verified using nucleotide BLAST (Johnson et al., 2008). Based on Galisa et al. (2012) work, recA, gyrB and rho were selected as the reference genes candidates and the qRT PCR primers were designed as previously mentioned (Galisa et al., 2012).

The amplification efficiencies of qRT primers were evaluated using the genomic DNA (gDNA) of WT *K. xylinus*. The gDNA was isolated from late exponential phase cells using GeneJet Genomic DNA Purification Kit (Thermo Fischer Scientific, USA), as per the manufacturer’s instructions. The qRT PCR mix were prepared in PowerTrack SYBR Green Master Mix (Applied Biosystems, USA). The target genes were amplified from 8 ng of gDNA template with CFX96 Touch Real-Time PCR (Bio-Rad, USA) under the following conditions: initial denaturation at 95°C for 2 minutes, followed by 40 cycles of 95°C for 15 seconds (denaturation) and 60°C for 1 minute (annealing and extension), with a plate read step. For the dissociation phase, the protocol included 95°C for 15 seconds, followed by melt curve analysis from 60°C to 95°C with a 0.5°C increment per minute, and concluded with a plate read step.

Recombinant *K. xylinus* cells, harbouring the post-transcriptional regulation cassette that contains the sRNA sequence targeting bcsD, bcsZ and ccpA mRNA, were pre-cultivated in HS medium containing 2% cellulase, 20 g L^-1^ glucose and 340μg ml^-1^ Cm at 30°C and 180 rpm. After 2 days of cultivation, three biological replicates for each inducer concentration (100 nM and 500 nM) were reinoculated (final OD600nm, 0.1) into the same medium containing appropriate AHL concentrations and grown at 30°C and 180 rpm. Subsequently, the total RNA was extracted using GeneJET RNA Purification Kit (Thermo Fischer Scientific, USA) and analyzed using Qbit fluorometric quantification (Thermo Fischer Scientific, USA) using the Qbit RNA HS standard. Template RNA of 2.5 ug was used for cDNA synthesis using 5X Superscript IV ViLo Master mix (Thermo Fischer Scientific, USA), and analyzed using Qbit. For qRT PCR, the cDNA of both control (noninduced) and treated (AHL induced) samples were amplified (in triplicates) with QrecA (reference gene) and respective gene-specific qRT PCR primers (Table 1), using the above-mentioned reaction conditions. The fold change in relative gene expression was calculated using 2^−ΔΔ*C*^_T_ method (Livak & Schmittgen, 2001).

### BC production and pretreatment

To study the effect of post-transcriptional regulation of the target genes on BC synthesis, both WT and recombinant *K. xylinus* cells were statically cultivated in HS medium containing 20 g L-^1^ glucose. For recombinant strains, Cm (340μg ml^-1^) and appropriate AHL concentrations were supplemented to the growth medium. The cells were statically grown in 30°C for 5 days. Following BC production, the pellicles were subjected to alkali treatments and washed thoroughly to inactivate the entrapped cells and remove medium components, respectively, as described in Mangayil et al. (2017) (Mangayil et al., 2017). Subsequently, the washed BC pellicles were oven-dried overnight at 60°C on pre-weighed weighing boats and the dry weight was calculated.

### Analytics

Cell biomass was determined by measuring the OD_600nm_ values using a spectrophotometer (Ultraspec 500pro, Amersham Biosciences, UK). Presence of gluconate in WT *K. xylinus* and *K. xylinus* Δ*gdh* were analyzed using a high-performance liquid chromatography (HPLC) equipped with Shodex SUGAR column (300 mm × 8 mm, Phenomenex), autosampler (SIL-20AC HT, Shimadzu), refractive index detector (RID-10A, Shimadzu), and 0.01 M H_2_SO_4_ as the mobile phase. BC dry weight measurements were conducted using an analytical balance (ES 220A, Precisa, Switzerland).

The X-ray diffraction patterns of the dried BC films were analyzed using a X-ray diffractometer (XRD; Empyrean multipurpose diffractometer, PANalytical B.V, US) and the crsystaline indices were calculated using peak deconvolution method as previously mentioned in Mangayil et al. (2021) (Mangayil et al., 2020; Park et al., 2010). Wide-angle X-ray scattering (WAXS) analysis of the oven dried BC sheets were conducted as described in Penttilä et al. (2020) (Penttilä et al., 2020).

## Results and Discussion

### Investigating the suitability of an sRNA-based post-transcriptional repression system in *K*. *xylinus* using genome-integrated *mRFP1*

This study aims in investigating the effect of post transcriptional regulation of *bcsD, ccpA* and *bcsZ* genes on BC crystallinity. Florea et al. (2016) have reported a gene repression cassette for *K. rhaeticus* iGEM strain (Florea et al., 2016). To test whether such system would be suitable for *K. xylinus*, a genome-integrated *mRFP1* strain, targeting to replace the *gdh* gene, was constructed. Based on the reports by Shigematsu et al. (2005) and Kuo et al. (2015), the quinoprotein glucose dehydrogenase at the genome location 814017 – 816407 bp was targeted for homologous recombination (C.-H. Kuo et al., 2015; Shigematsu et al., 2005). After visual inspection, colony PCR and confirmed constitutive mRFP1 expression (from gLF-JmRFP1-CmR-gRF cassette in pSEVA241) in *E. coli* XL1, the plasmid was subsequently transformed into *K. xylinus*. Visual inspection of mRFP1 gene expression from transformants (Supplementary Fig. S1) and BC pellicles (Supplementary Fig. 2), and absence of gluconate from HPLC from the cultivation samples confirmed the deletion of *gdh* gene. Subsequently, the mRFP1 expression from *K. xylinus* Δ*gdh* (*Kx*Δ*gdh*) was compared with *K. xylinus* harboring J23104-mRFP plasmid. The relative fluorescence values from plasmid-based expression were approximately five times higher than those from genomic expression. This lower gene expression in genome-integrated cells was anticipated due to their single-copy configuration.

Next, we transformed *Kx*Δ*gdh* with the sRNA construct, JsRFPk, and investigated cell growth and mRFP1 gene repression. While the non-induced *Kx*Δ*gdh*-JsRFPk constructs showed robust growth, reaching a final OD_600nm_ of 4.8±0.3, severe growth inhibition was observed in AHL induced cultivations (Figure 1). After 24 hours of growth, a 56% reduction in OD_600nm_ was observed in cultures induced with 10 nM AHL, with this reduction increasing to 84% at higher inducer concentrations (Figure 1A). Over 48 hours, the OD_600nm_ drop intensified to 95%, followed by a slight recovery in growth at 72 hours. By 144 hours, growth inhibition was alleviated, with the cells reaching a final OD_600nm_ of 5.2 – 5.9, except for those cultivated in 10 nM AHL.

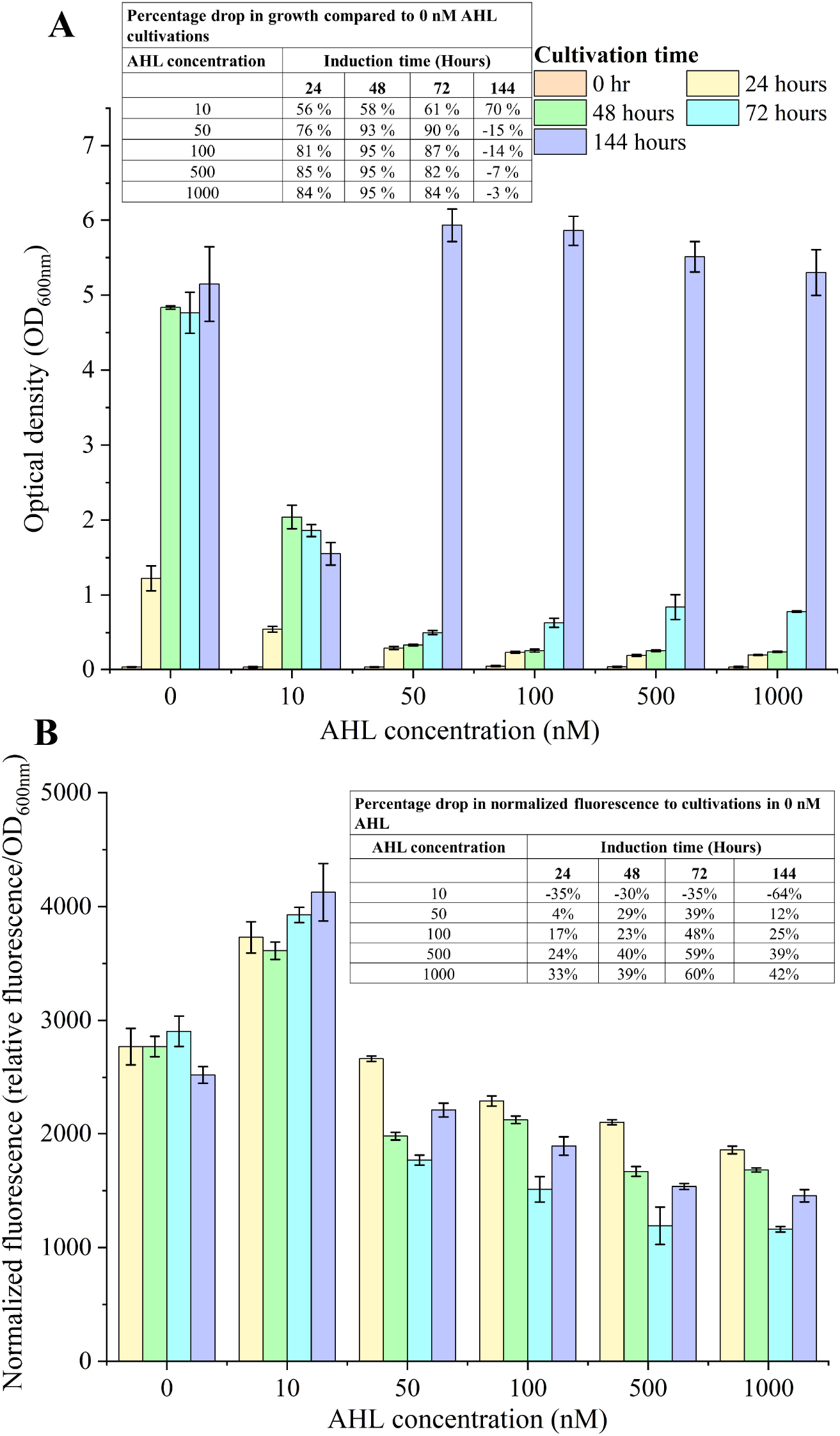

As observed from the normalized fluorescence data, cultures induced with 10 nM AHL did not repress mRFP1 expression. Nevertheless, with increased AHL concentrations there was a consistent trend of increased repression in mRFP1 gene expression (Figure 1B). However, after 144 hours of cultivation, the percentage drop in fluorescence decreased. These variations in the normalized fluorescence data are likely due to the cellular stress experienced during the first three days of cultivation. As shown in Figure 1A, cells gradually adapted to the stress, leading to a steady increase in OD_600nm_ and ultimately reaching the growth levels similar to those of non-induced cells by 144 hours. At this time point, mRFP1 expression was repressed by 12%, 25%, 39%, and 42% at AHL concentrations of 50 nM, 100 nM, 500 nM, and 1000 nM, respectively. These tests indicate that despite the impact on growth, the post-transcriptional repression of genome-integrated mRFP1 was effective.

We reason that the overexpressed *E. coli* Hfq could be the reason for the observed growth inhibition. Although such effects were not investigated in *K. rhaeticus* iGEM (Florea et al., 2016), growth inhibitions due to overexpressed Hfq chaperone have been reported for several Gram-positive and Gram-negative bacteria (Fernández et al., 2016; Keefer et al., 2017). Since the Hfq RNA chaperone is endogenous in *K. xylinus* (Genomic location, 2818175 – 2818453 bps), we tested whether the observed growth inhibition could be alleviated by removing the Hfq expression operon (pLux01-RBS-Hfq) from JsRFPk (Park et al., 2021). The *Kx*Δ*gdh*-JsRFPΔHfqk cultivations showed a significant reduction in growth inhibition, decreasing from 56%–95% in *Kx*Δ*gdh*-JsRFPk (Figure 1A, Inset Table) to 4%–23% in *Kx*Δ*gdh*-JsRFPΔHfqk (Figure 2A, Inset Table). The role of Hfq, a global RNA chaperone, differs significantly between its endogenous and overexpressed forms. Endogenous Hfq, naturally present in bacteria, is essential for post-transcriptional regulation, facilitating interactions between sRNAs and their target mRNAs, which are crucial for stress responses, virulence, and adaptation to environmental changes (Dutta & Srivastava, 2018; Morita & Aiba, 2019). However, overexpression of non-native Hfq can disrupt the balance of sRNA-mRNA interactions, resulting in gene regulation and growth inhibition due to excessive interference with native regulatory networks or reducing their availability for normal regulatory functions (Cech et al., 2016; Gottesman & Storz, 2011). In terms of RFP expression, normalized fluorescence observed from *Kx*Δ*gdh*-JsRFPΔHfqk (13% -43%; Figure 2B) was comparable to that seen in *Kx*Δ*gdh*-JsRFP cultivations (4% -60%; Figure 1B). These data suggests that removal of Hfq expression operon effectively repressed post-transcriptional repression while reducing the metabolic burden. This indicates that a repression plasmid lacking the Hfq expression operon is an ideal construct for *Komagataeibacter* spp, and thus was employed to study post-transcriptional repression of bcsD, bcsZ and ccpA.

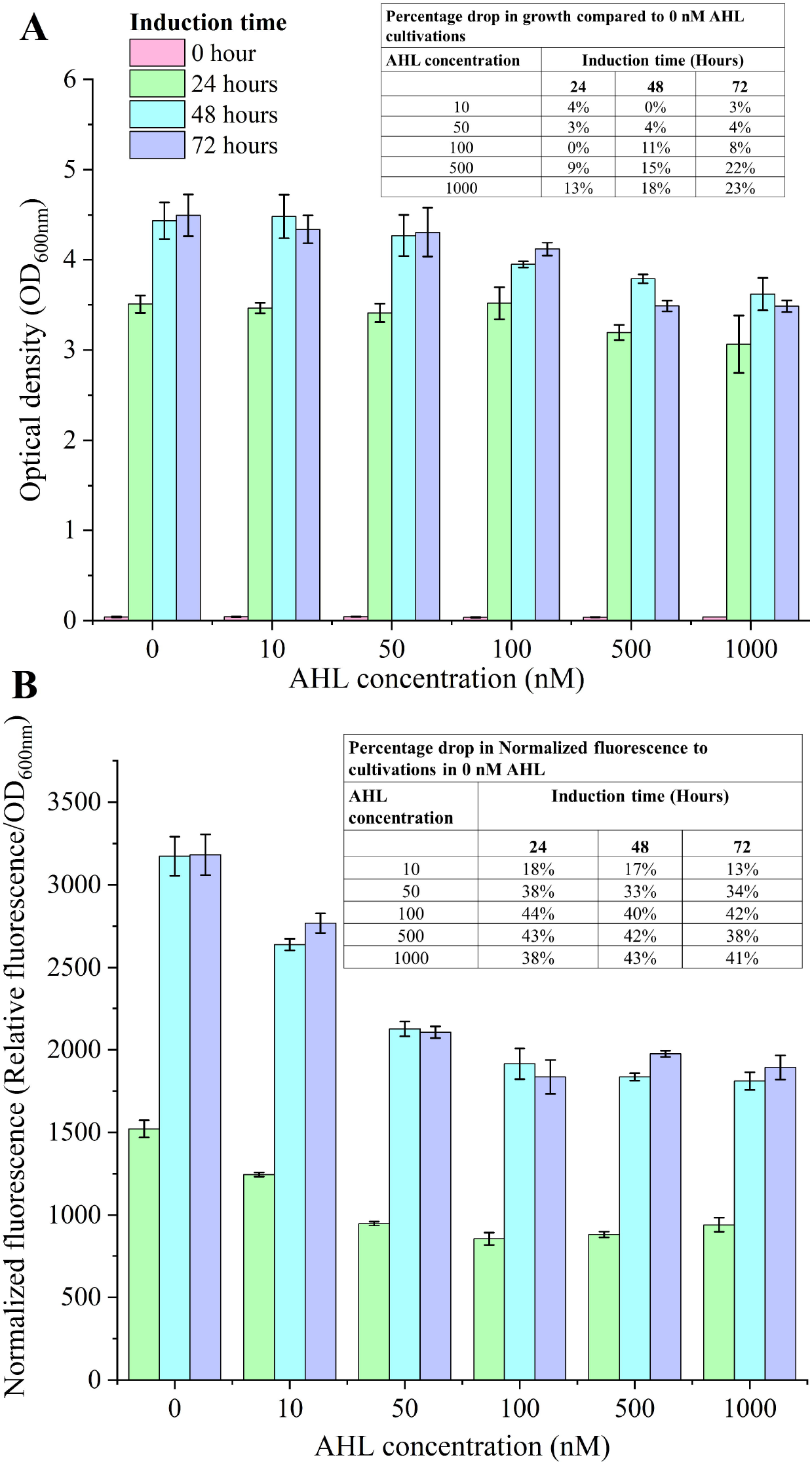

### sRNA-based post transcriptional repression of *bcsD, bcsZ* and *ccpA*

Before conducting the qRT PCR experiments, the amplification efficiencies of the qRT primers towards the target (*bcsD, bcsZ* and *ccpA*) and reference housekeeping (*recA, rho* and *gyrB*) genes were assessed using *K. xylinus* gDNA (Galisa et al., 2012). Analysis of the amplification and melt peak curves demonstrated that the qRT primers were specific to the target and *recA* and *rho* reference genes (Supplementary figures S4 – S6). However, since the melt peak data for *gyrB* revealed two peaks, indicating the presence of two amplicons, it was excluded from further experiments. To identify the optimal reference gene for qRT experiments, the average Cq values from the *recA* and *rho* qRT PCR data were analyzed using the RefFinder tool (https://www.ciidirsinaloa.com.mx/RefFinder-master) (Xie et al., 2023). The analysis of Cq values from quadruplet replicates indicated *recA* as the most stable reference gene (Supplementary Fig. S7).

Based on the binding energy predictions (Table 2) and the observed repression of mRFP1 expressed under a strong synthetic constitutive promoter, we hypothesized that AHL-induced transcription of sRNAs designed to specifically target the genes in this study could lead to significant post-transcriptional repression. Reflecting the mRFP repression data (Figure 2), we tested the effect of 0 nM (control), 10 nM and 100 nM AHL concentrations on post-transcriptional regulation of bcsD, bcsZ and ccpA.

Initial qRT-PCR experiments with *Kx*Δ*gdh*-JsbcsD24ΔHfqk cells (carrying a repression plasmid with a 24-bp sRNA targeting bcsD mRNA) induced with 10 nM and 100 nM AHL showed low levels of repression compared to the genome-integrated mRFP. The fold change measured by qRT-PCR, revealed that 10 nM AHL was insufficient to trigger post-transcriptional repression of *bcsD* and cultures induced with 100 nM AHL showed only 0.6% decrease in gene expression. One possible explanation could be the timing of mRNA sampling, as the total mRNA samples were collected from cells that had grown for 5 days and reached an OD_600nm_ of 1.0–1.2. Subsequent experiments were designed to sample the total mRNA from cells at the exponential growth phase (OD_600nm_ 0.6–0.7). Additionally, effect of higher inducer concentration (500 nM) on post-transcriptional repression was also tested.

Using cDNA from exponentially grown cells induced with 100 nM and 500 nM AHL, we observed a correlation between inducer concentration and gene repression. The qRT-PCR results from *K. xylinus* cells carrying the repression cassettes with 24-25 bp sRNA sequences showed 28% -34%, 11% -24%, and 4.4% -20% repression of bcsD, bcsZ, and ccpA transcription, respectively. As anticipated, constructs with 12-bp sRNA sequences exhibited lower levels of gene repression, with more pronounced effects at higher inducer concentrations.

### Effect of post transcriptional repression of bcsD, bcsZ and ccpA on BC production

Next, we aimed to evaluate the impact of repressing individual target genes on BC production and structural variations. Since the qRT-PCR data from *Kx*Δ*gdh*_JsbcsZ12ΔHfq and *Kx*Δ*gdh*_JsccpA12ΔHfq cells induced with 100nM AHL did not indicate transcriptional repression, these samples were excluded. The mean BC titers values from WT and recombinant *K. xylinus* cells harboring the sRNA-based post-transcriptional repression plasmid, cultivated in the presence of 100 nM and 500 nM AHL are presented in Figure 3. The tested samples produced BC titers ranging from 1.5 – 2.7 g L^-1^. As seen in the BC titers from WT cells (2.5±0.3 g L^-1^ devoid of inducer, 2.4±0.3 g L^-1^ with 100 nM AHL, and 2.7±0.2 g L^-1^ with 500 nM AHL), we can observe that the inducer did not negatively affect BC production capacity.

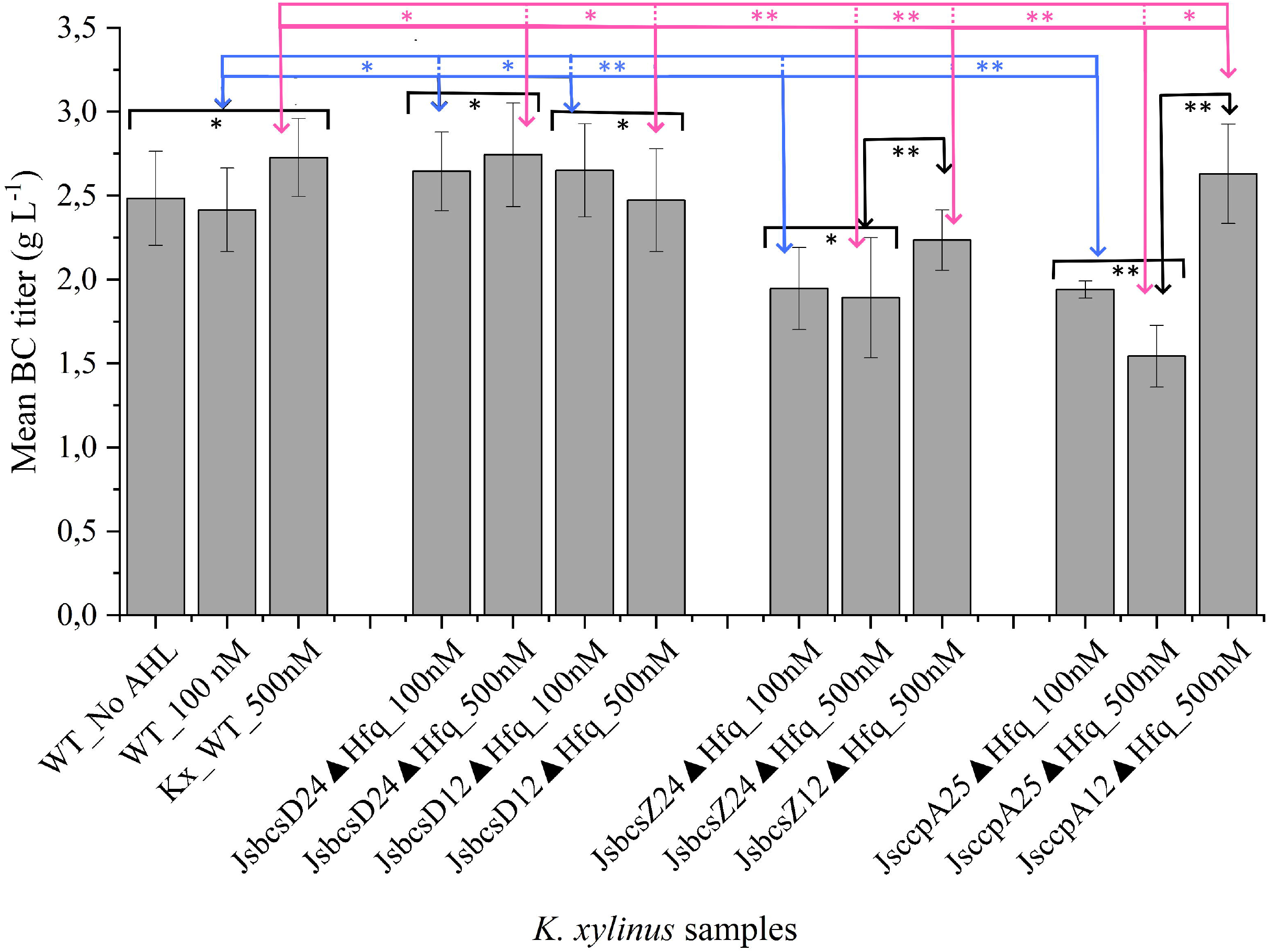

bcsD, a component of the bcs terminal complex, interacts with bcsAB in nanofibril assembly, assists bcsC in nanofibril extrusion, and interacts with accessory proteins (Kondo et al., 2022; Mehta et al., 2015; Sana et al., 2024; Saxena et al., 1994; Sunagawa et al., 2013). In addition to its involvement in the hierarchical assembly of the terminal complex, bcsD contributes to the crystalline characteristics of nascent nanofibrils. Based on previous bcsD knockout and overexpression studies, one might expect that post-transcriptional repression of bcsD would result in a significant reduction in BC production (Mangayil et al., 2017; Yang et al., 2024). However, our experiments did not confirm this expectation. Although qRT-PCR data showed a ∼1.2 – 1.8-fold change (in cells with 24-bp and 12-bp sRNA sequences, respectively) in relative gene expression, BC titers remained consistent, with 2.7 g L^-1^ and 2.4 – 2.7 g L^-1^ produced from cultures induced with 100 nM and 500 nM AHL, respectively. We speculate that while the gene was repressed, the 28% to 34% repression observed in the qRT-PCR analysis was insufficient to significantly impact BC production.

However, in *K. xylinus* cells with repression plasmids targeting the *bcsZ* and *ccpA* genes, a more pronounced and statically significant difference in BC titers were observed. This observation aligns with the qRT-PCR results (Table 3). For example, the qRT-PCR data from *K. xylinus* cells with repression plasmids containing 24-bp sRNA sequences showed ∼2.8-fold reduction in relative bcsZ gene expression in cultures grown with 100 nM and 500 nM inducer. Gene knockout studies and the exogenous addition of endoglucanases have demonstrated that the *bcsZ* gene significantly impacts BC production (Nakai et al., 2013; Tonouchi et al., 1995). We also observed a significant drop in BC titer with tighter post-transcriptional repression (Figure 3). Cells containing the repression plasmid with a 24-bp sRNA sequence, when induced with 100 nM and 500 nM AHL, showed ∼19.5% and ∼30% reductions in BC titer, respectively, compared to WT cells grown under the same inducer concentrations. Nevertheless, within the recombinant cells induced with same Ahl concentration, the BC titers remained consistent, indicating that 100 nM AHL is sufficient to trigger post-transcriptional repression. As hypothesized when designing the sRNA constructs, cells with the shorter 12-bp sRNA sequence exhibited milder post-transcriptional repression (Table 3), thereby resulting in higher BC titers (2.3 g L^-1^) compared to cells with the 24-bp sRNA sequence (Figure 3).

**Table 3.** qRT PCR results.

| Table 3. qRT PCR results |  |  |
| --- | --- | --- |
| Target | Decrease in relative gene expression with respect to uninduced cultivation |  |
|  | 100 nM AHL | 500 nM AHL |
| *24-25 bps of sRNA complementary to the target mRNA* |  |  |
| *bcsD* | 28% | 34% |
| *bcsZ* | 11% | 24% |
| *ccpA* | 4.4% | 20% |
| *12-bps of sRNA complementary to the target mRNA* |  |  |
| *bcsD* | 6% | 11% |
| *bcsZ* | - | 8.7% |
| *ccpA* | - | 2.3% |

The *ccpA* gene, located downstream of *bcsZ* in the bcs operon, is known to be crucial for the localizing the bcs terminal complex through interactions with bcsD (Kondo et al., 2022; Sana et al., 2024; Standal et al., 1994; Sunagawa et al., 2013). This study observed negative effects on BC titers due to post-transcriptional repression of ccpA. Specifically, cells with the repression plasmid carrying 25 bps sRNA sequence showed a more pronounced reduction in BC titers. The *Kx*Δ*gdh*_JsccpA25ΔHfq cells experienced a statistically significant reduction in BC titers when induced with either concentration of the tested inducers, with a more substantial drop at higher AHL concentrations. In contrast, the BC titers from *Kx*Δ*gdh*_JsccpA12ΔHfq cells, which had a 2.3% reduction in relative gene expression, were comparable to those from WT cultures under similar induction conditions. This indicates that, similar to the effect seen with *bcsZ*, using a shorter sRNA sequence can mitigate the severe impact of ccpA repression on BC production.

The BC synthesized by both WT and recombinant *K. xylinus* cells was subsequently subjected to structural characterization using XRD, and WAXS. Surprisingly, the XRD analysis revealed that all tested BC samples had relatively high crystallinity index (CI) values (>97%), which are higher than those typically reported in the literature. However, variations in the amorphous content were observed (Figure 4). Among the BC synthesized by cells induced with 100 nM AHL, only *Kx*Δ*gdh*_JsbcsD24ΔHfq exhibited a significant variation in amorphous content compared to the BC produced by WT cells under similar conditions. Pronounced variations were observed in BC synthesized by recombinant cells induced with higher inducer concentrations, indicating that the repression cassette effectively modulated BC crystallinity.

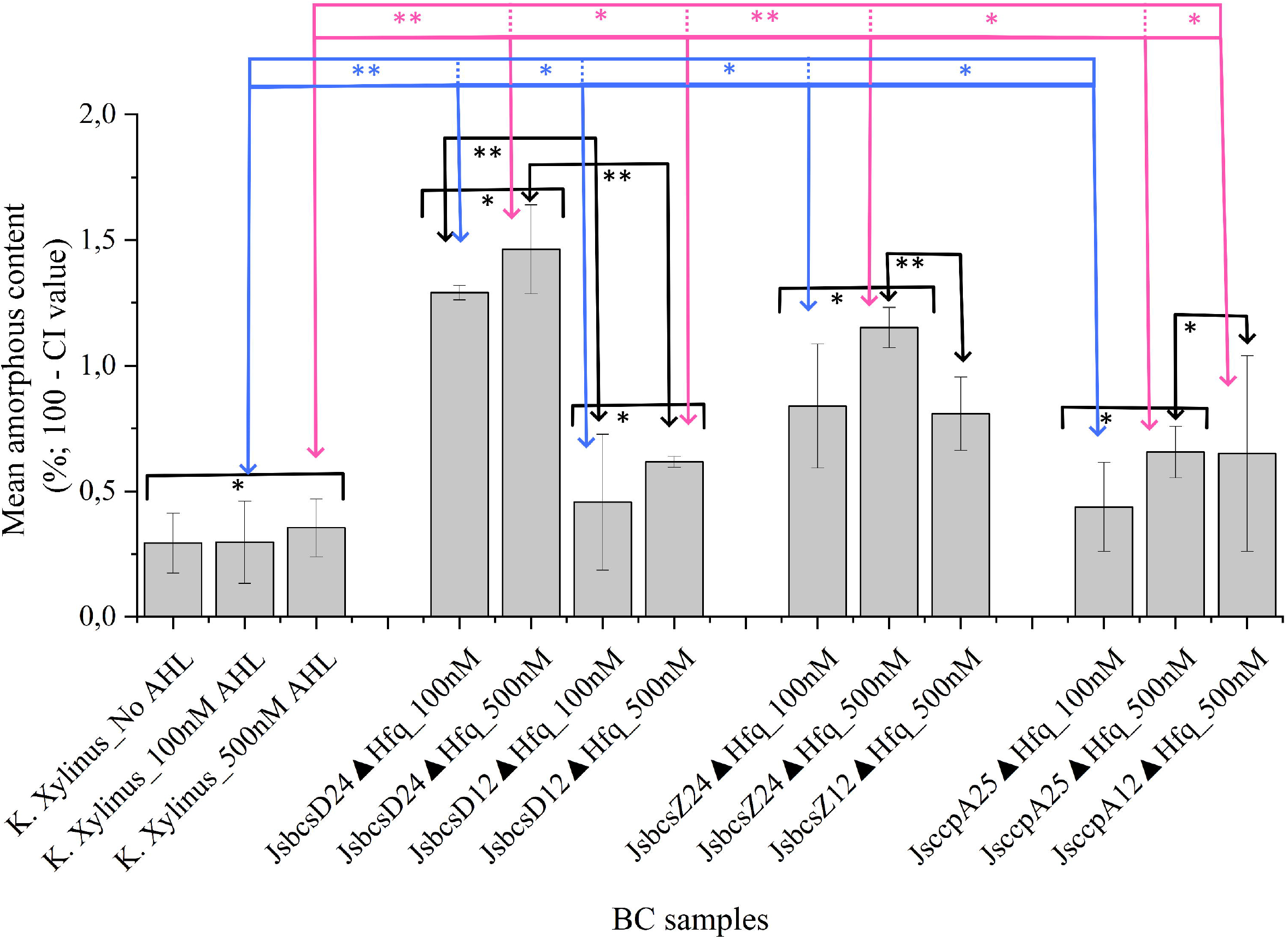

To further validate the structural changes observed from the XRD data, the BC samples were subjected to WAXS (Figure 5). The WAXS data of the tested BC films indicated the major cellulose I diffraction peaks at ∼14-17° (equatorial peak), ∼21°, ∼22° and ∼35° representing the characteristic 1-10, 110, 102, 200 and 004 reflections of triclinic cellulose Iα crystal form (Chiriac et al., 2014; Rovera et al., 2022). Literature reports that in the event of an undisturbed cellulose I crystallite structure, the first predominant peak from a WAXS diffraction profile will be a broad peak that combines the 110 and 1-10 reflections (Chiriac et al., 2014; Jungnikl et al., 2008; Lutz-Bueno et al., 2022). However, with disturbances to the crystallinity, the peaks diverge, forming two evident peaks as observed from this study (Chiriac et al., 2014; Ibbett et al., 2013). As identified by Rovera et al. (2021), the emergence of additional peaks at ∼29° and ∼32.5° could be caused by the incomplete removal of ammonium hydroxide crystals used in the pretreatment of BC films (Rovera et al., 2022).

We observed broad peaks at ∼28° -30° and ∼30°-32°. The peak at ∼28°-30°, that showed no obvious profile, could be attributed to the presence of NaOH crystals from BC pretreatments. However, much sharper peak at ∼30°-32° that shows a distinct profile could signify the amorphous phase in the BC microstructure (Lutz-Bueno et al., 2022). Additionally, the peak profile at ∼30°-32° corelates with the amorphous content calculated from the XRD data. For instance, the samples that showed statistically significant values in the amorphous content, i.e., BC synthesized from *Kx*Δ*gdh*_JsbcsD24ΔHfq, *Kx*Δ*gdh*_JsbcsZ24ΔHfq and *Kx*Δ*gdh*_JsccpAD24ΔHfq induced with 500 nM AHL (Figure 4) demonstrates high peaks at ∼30°-32°. These may suggest that the alterations in BC crystallinity was successfully induced through post-transcriptional repression.

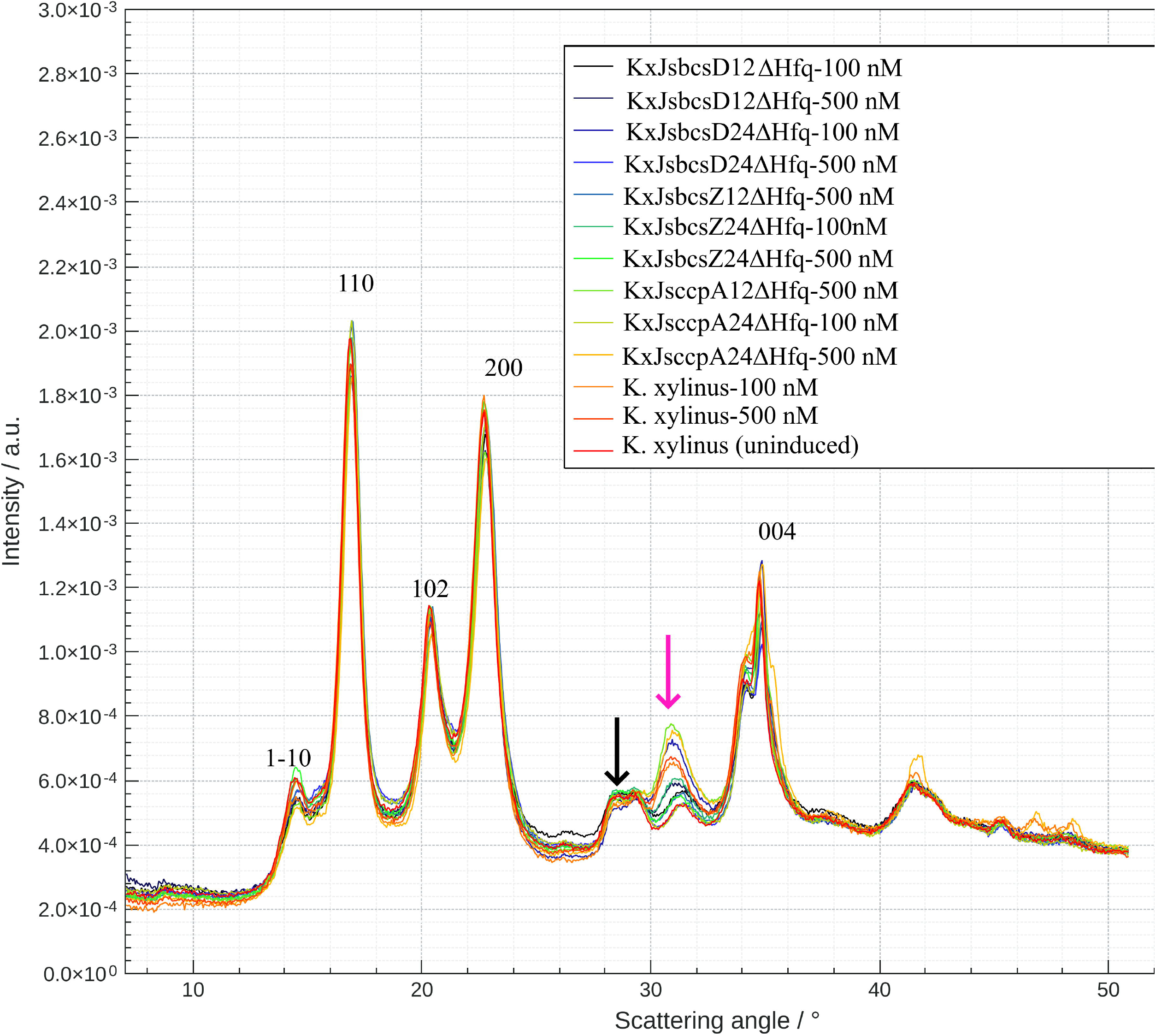

## Conclusion

In summary, this study aimed to understand the impact of post-transcriptional repression of the bcsD, bcsZ, and ccpA genes on BC crystallinity. During the construction of the repression cassette, we discovered that overexpressing the Hfq RNA chaperone imposed a significant metabolic burden on *K. xylinus*. This burden was reduced by removing the RNA chaperone, leading to 18% to 43% post-transcriptional repression of the genome-integrated mRFP1 gene. By designing repression constructs with sRNA sequences of 24-25 and 12 bases complementary to the target mRNA, and by inducing with different inducer concentrations, we successfully achieved varying levels of repression in *bcsD, bcsZ*, and *ccpA* gene expression. Although BC production remained largely unchanged in bcsD-repressed cells, statistically significant reductions in BC titers were observed in cells with repression cassettes targeting bcsZ and ccpA. These effects were more pronounced at higher inducer concentrations and in cells with repression cassettes containing 24-25 base sRNA sequences. Furthermore, the variations in the amorphous content of BC samples identified through XRD and WAXS diffraction patterns correlated with the post-transcriptional repression data. Although this study successfully demonstrated the effects of individual gene repression on BC titers and structural variations, it also highlights the need for further engineering to fine-tune the repression cassette. Future research may focus on developing a cassette for the combined post-transcriptional repression of the genes targeted in this study, aiming to systematically control BC crystallinity without compromising the titers.

## Supporting information

File containing supllementary figures

## CRediT authorship contribution statement

R.M. conceptualized the work, designed, and conducted the experiments, interpreted the data, wrote the original draft and acquired funding. E.S. conducted the XRD analysis. T.E. and V.S. supervised the work and reviewed the manuscript.

## Acknowledgments

This study was funded by the Research Council of Finland (Grant numbers, 346983 and 353673 for R.M.).

## Appendix Appendix A. Supplementary data

The manuscript is accompanied with a supplementary file containing supplementary figures.

## Data availability

All the data used in this study are presented within the manuscript text or in the supplementary file.

