## Supplementary material for "Modulating Bacterial Nanocellulose Crystallinity through Post-Transcriptional Repression in *Komagataeibacter xylinus*": File containing supllementary figures

### Supplementary figures

A B

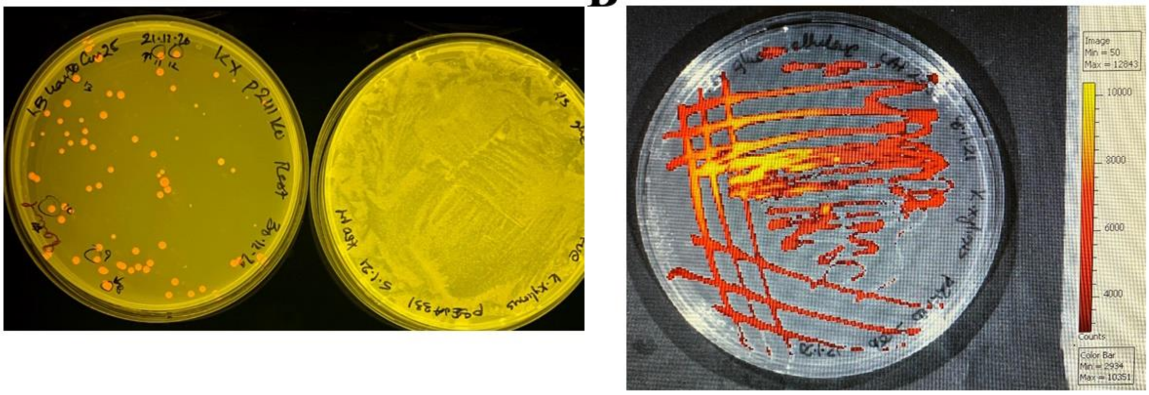


Supplementary Figure S1. Visual inspection of *K. xylinus*Δgdh transformants with (A) genome integrated J23104-mRFP1-CmR and non-transformed cells under blue light and (B) under IVIS optical imaging station.

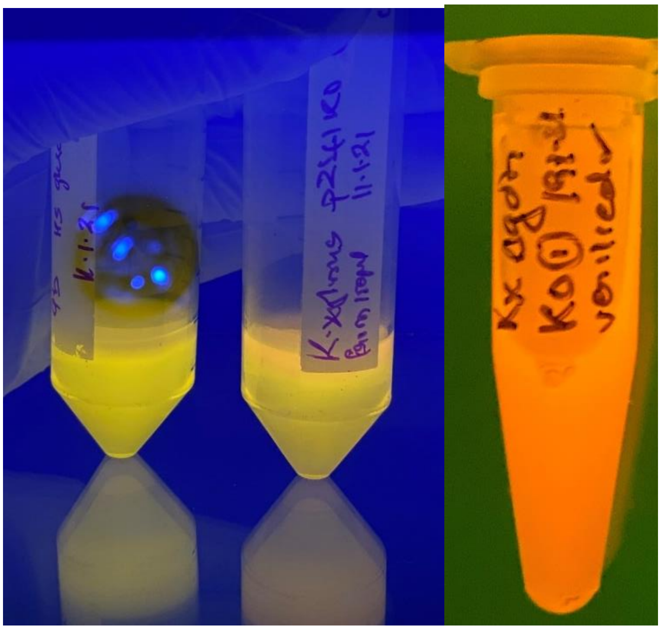

Supplementary Figure S2. mRFP1 expression in BC pellicles synthesized by WT (left) and *K. xylinus* Δ*gdh* (right) cells.

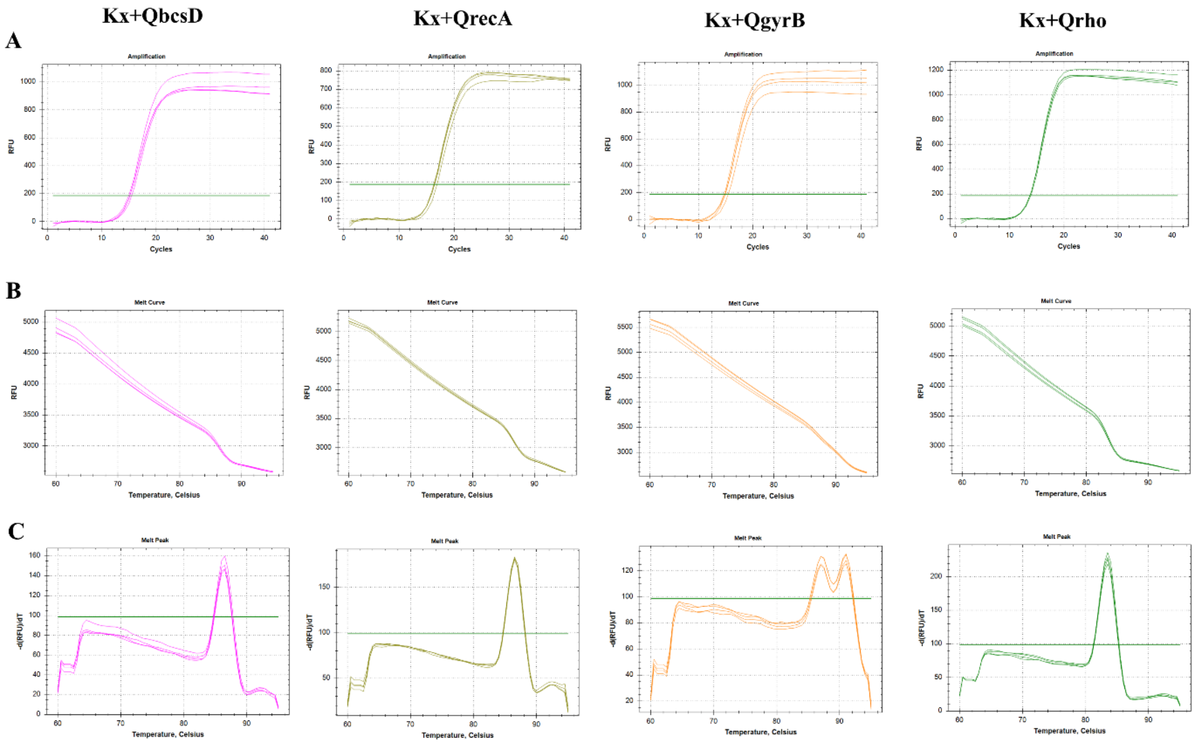


Supplementary figure 4. Preliminary data indicating the specificity of the designed qRT primers towards the target genes (*bcsD*, *recA*, *gyrB* and *rho*). The figures demonstrate the (A) amplification, (B) melt curve and (C) melt peaks data.

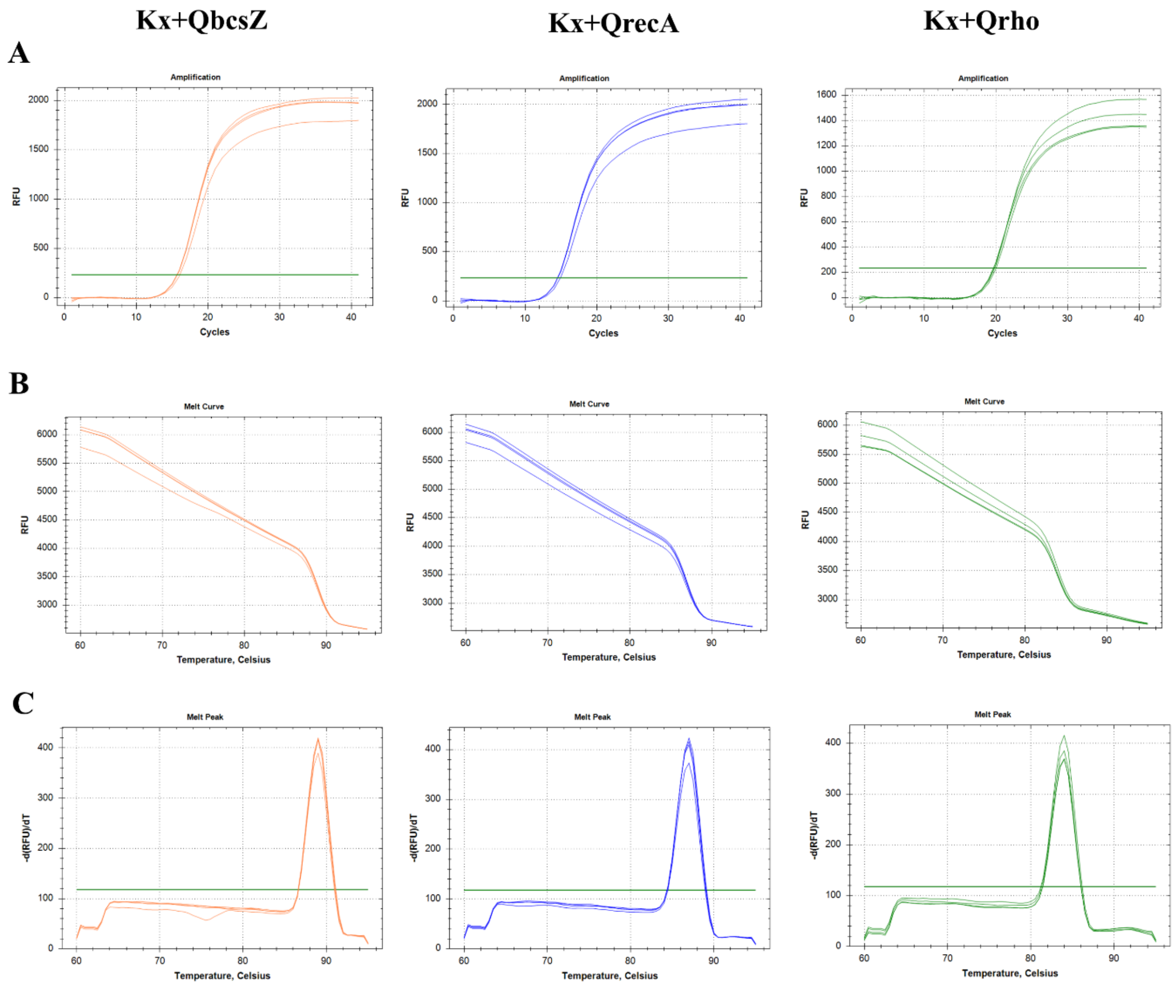


Supplementary figure 5. Preliminary data indicating the specificity of the designed qRT primers towards the target genes (*bcsZ*, *recA* and *rho*). The figures demonstrate the (A) amplification, (B) melt curve and (C) melt peaks data.

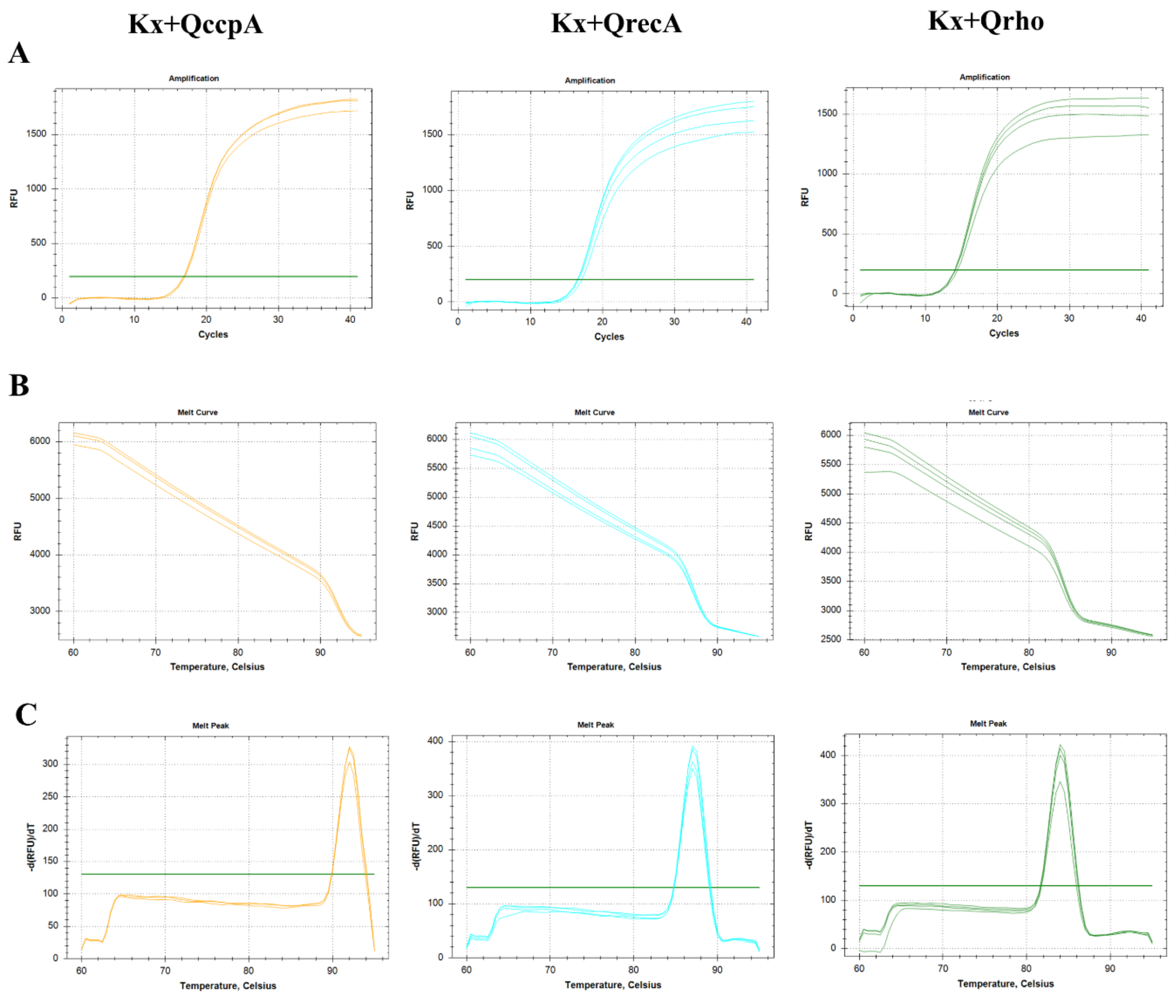


Supplementary figure 6. Preliminary data indicating the specificity of the designed qRT primers towards the target genes (*ccpA*, *recA* and *rho*). The figures demonstrate the (A) amplification, (B) melt curve and (C) melt peaks data.

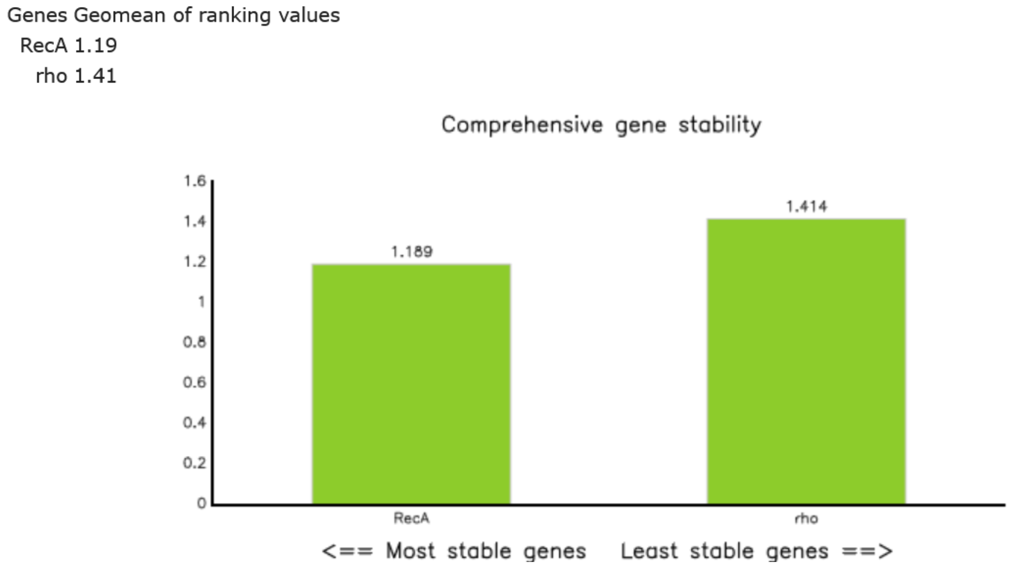


Supplementary figure 7. Gene stability analysis of reference housekeeping genes using the RefFinder tool (https://www.ciidirsinaloa.com.mx/RefFinder-master) (Xie, Wang, & Zhang, 2023)
